# Compensation masks trophic cascades in complex food webs

**DOI:** 10.1101/068569

**Authors:** Ashkaan K Fahimipour, Kurt E Anderson, Richard J Williams

## Abstract

Ecological networks, or food webs, describe the feeding relationships between interacting species within an ecosystem. Understanding how the complexity of these networks influences their response to changing top-down control is a central challenge in ecology. Here, we provide a model-based investigation of *trophic cascades* ‒ an oft-studied ecological phenomenon that occurs when changes in the biomass of top predators indirectly effect changes in the biomass of primary producers ‒ in complex food webs that are representative of the structure of real ecosystems. Our results reveal that strong cascades occur primarily in low richness and weakly connected food webs, a result in agreement with some prior predictions. The primary mechanism underlying weak or absent cascades was a strong compensatory response; in most webs predators induced large population level cascades that were masked by changes in the opposite direction by other species in the same trophic guild. Thus, the search for a general theory of trophic cascades in food webs should focus on uncovering features of real ecosystems that promote biomass compensation within functional guilds or trophic levels.

## 1 Introduction

Trophic cascades occur when changes in an ecosystem's top trophic level propagate down through the food web and drive changes in the biomass of primary producers (Hairston et al, 1960; Paine, 1980). Cascades have now been documented in virtually every type of ecosystem, but neither conceptual nor mathematical theories have been able to explain widespread variation in observed cascade strengths (Borer et al, 2005; Shurin et al, 2010); in some ecosystems, strong cascades impact several lower trophic levels while in others they diminish within a single trophic level (Heath et al, 2014). Indeed, *trophic trickles* – weak or absent cascades in response to major changes to a food web's top trophic level — abound in nature (McCann et al, 1998; Mikola and Setala, 1998; Halaj and Wise, 2001). Given that human actions are disproportionately altering biomass of top predators (Estes et al, 2011), there is a pressing need to understand under what circumstances such changes will or won’t cascade through complex food webs (Terborgh et al, 2010).

Food web structure has long been predicted to regulate cascade strength (Strong, 1992; Pace et al, 1999; Polis et al, 2000; Shurin et al, 2010) and the magnitudes of indirect effects in general (MacArthur, 1955; Yodzis, 2000). Indirect tests of this hypothesis have so far been accomplished by leveraging data on community features like functional or taxonomic diversity (Borer et al, 2005; Frank et al, 2006), in hopes that these proxies for web structure could provide clues to the features of ecological networks that influence the magnitude of cascading top down effects. However results have been mixed, with studies reporting both strong (Frank et al, 2006, 2007; Baum and Worm, 2009) and weak or noisy (Borer et al, 2005; Fox, 2007) associations between diversity measures and cascade strengths. Whether data support assertions that food web structure regulates cascade strengths remains unclear, and a coherent understanding of when relatively strong or weak trophic cascades occur is still lacking.

One impediment to progress is that extensions of cascade theory toward species rich and topologically complex food webs are needed to guide further empirical study. To date, cascade theory has focused largely on understanding variation in cascade strengths in model food chains (Oksanen et al, 1981; McCann et al, 1998; Heath et al, 2014; DeLong et al, 2015) and although extensions of cascade theory to alternate trophic modules exist (Bascompte et al, 2005; Fahimipour and Anderson, 2015), the mechanisms underlying variation in cascade strength in species rich and complex trophic networks remain poorly understood (Holt et al, 2010; Shurin et al, 2010).

Here we use a bioenergetic food web model to explicitly study the emergence of trophic cascades in species rich webs that are representative of the structure of real ecosystems following the invasion of a novel top generalist predator. We demonstrate that the strongest trophic cascades occur in small and weakly connected food webs – a result in agreement with some prior predictions (Pace et al, 1999; Polis et al, 2000; Fox, 2007; Shurin et al, 2010). Moreover, our results reveal that biomass compensation within producer and consumer functional guilds, whereby some species increase in biomass while others decrease proportionately, is the most common mechanism underlying weak or absent trophic cascades. Thus, the search for a general theory of trophic cascades in food webs should focus on uncovering the abiotic and biotic features of real ecosystems that promote or preclude biomass compensation and compensatory dynamics within functional guilds.

## 2 Methods

We implemented a modeling framework similar to that described by Yodzis and Innes (1992) and reviewed by Williams et al (2007). Namely, we generated multitrophic level food web topologies using an ecological niche model (Williams and Martinez, 2000) and simulated the dynamics of energy flows on these generated webs using a bioenergetic model (Yodzis and Innes, 1992; Brown et al, 2004; Brose et al, 2006b; Williams et al, 2007). This modeling framework was chosen because it is grounded in empirical knowledge about network structure, species parameters and nonlinear interaction dynamics. Previous work has shown that allometric scaling of parameters and complex functional responses are vital for modeling persistent, complex multispecies food webs (Brose et al, 2006b; Boit et al, 2012), particularly when changes in species richness or web topology are imposed (e.g., Dunne and Williams, 2009). Because it is trivial to study cascades in model food webs that collapse upon predator invasion, we take advantage of previously studied features of this bioenergetic model (discussed below) to design more persistent systems that permit the study of cascades in the face of major changes to model web topology.

The niche model is discussed in detail by Williams and Martinez (2000), but briefly a one-dimensional niche axis on the interval [0, 1] is assumed and each species in the web is randomly assigned a “niche value” on this axis. Species *i* consumes all other species with niche values within a range on the axis, which is assigned using a beta function to randomly draw values from [0, 1]. This approach was used to generate realistic food web topologies (Williams and Martinez, 2000) for 1200 simulations in a factorial design: initial species richnesses of *S* = 10, 15, 20 and 25 were crossed with directed connectance *C* = 0.12, 16 and 0.2 as niche model parameters (4 richnesses × 3 connectances × 100 iterations = 1200 webs total). These values of *C* were chosen because they encompassed a wide range of empirically observed connectance values (Vermaat et al, 2009). Webs that deviated from the precise *C* values, contained disconnected nodes, or consisted of disconnected subgraphs were not considered.

Details of the energy flow model and parameters used herein are reviewed by Williams et al (2007). Namely, an allometrically scaled nonlinear bioenergetic model (Yodzis and Innes, 1992) was used to study the dynamics of species biomasses and the occurrence of trophic cascades in niche model webs when they are subject to the invasion of a new predator. We report results from a single ecologically reasonable set of model parameters, though similar results were obtained with other model parameterizations. Biomass dynamics were represented using the governing equations,

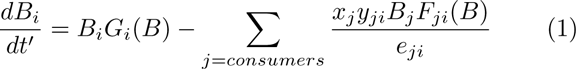

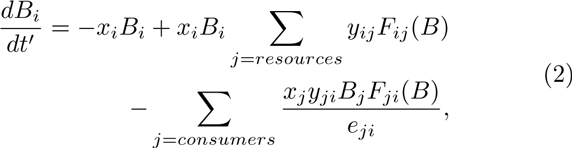

describing the dynamics of primary producers (e.g., plants; eq. 1) and consumers (e.g., herbivores, omnivores and higher trophic level predators; eq. 2). Here *B*_*i*_ is the biomass of species *i* and we use *R* and *N* when referring to producers or consumers respectively. All producers were assumed to have the same body mass, *M*_*r*_ = 1, and time *t’* was scaled with producer growth rate (see Williams et al, 2007 for details). To control for effects of varying productivity on trophic cascade strength, we maintained constant maximum productivity across simulations by assuming a systemwide carrying capacity *K* that is shared amongst *n*_*R*_ producer populations according to *K*_*i*_ = *K/n*_*R*_. Because of the well-documented effects of system productivity and enrichment on cascade strengths (e.g., Chase, 2003) we sought to constrain total potential productivity in all food webs, so that our results were not confounded by variation in the number of basal species generated by each niche model web.

In order to reduce the size of the parameter space being explored, all species in a web were assumed to have a constant consumer-resource body size ratio *Z*so that the mass of species *i* was *M*_*i*_ = *Z*^*P*^ where *P*is the length of the shortest path between species *i*and any producer at the base of the web. We report simulations in which *Z* = 42, so that for instance a linear three-species food chain comprising a producer, intermediate consumer and top predator would contain species with scaled body masses 1, 42 and 1764 respectively. This value of *Z* represents the mean predatorprey body mass ratio reported by Brose et al, 2006a, although the results presented herein were not sensitive to the choice of *Z* across its biologically relevant range.

The function *F*_*ji*_(*B*)is the normalized multi-species functional response for consumer *j* and resource *i*, developed by Yodzis and Innes, 1992 and extended by others (Brose et al, 2006b; Williams et al, 2007; Williams, 2008). To avoid the collapse of webs following predator invasions and permit the study of cascades after predator invasions, we explicitly considered a functional response that includes processes known to increase food web persistence in this model. These included the addition of mild interspecific consumer interference and slight relaxation of resource consumption when resources are very rare (Brose et al, 2006b). Adding consumer interference to a multispecies nonlinear functional can be represented as

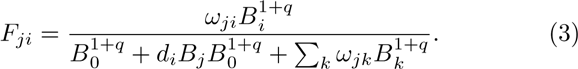

Here *d*^*i*^ is a positive constant that sets the amount of interference in the system and the sum in the denominator is over all *k* resources consumed by *j*. We assumed that interference was weak *(d*_*i*_ = 0.5) and set the shape parameter *q* = 0.2, which slightly relaxed consumption rates at very low resource biomasses – features that are well within the range of empirically observed functional responses (Brose et al, 2006b; Williams, 2008; Boit et al, 2012). We assume passive resource switching, so ω_ij_ = 1/*n*_*i*_ where *n*_*i*_ is the number of resources consumed by *j*.

Metabolic parameters in the bioenergetic model (Yodzis and Innes, 1992; Brose et al, 2006b) are given by

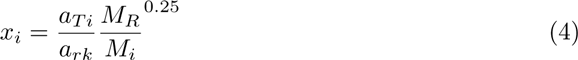

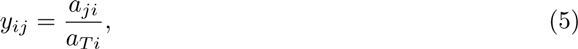

where *M*_*i*_ is the mass of an individual of species *i*and *M*_*r*_ is the mass of primary producers used for normalizing the time scale. The constants *a*_*T*_, *a*_r_ and *a*_*j*_ (*mass*^0.25^ × *time*^*-1*^) were previously determined from empirical data on the allometry of metabolism, production and maximum consumption respectively (Brose et al, 2006b). We assumed that all species were invertebrates, and so *a*_*r*_ = 1, *a*_*T*_ = 0.314 and *y*_*ij*_ = 8 (see Brose et al (2006b) for the derivation of these values). The metabolic parameter *x*_*i*_ is the mass specific metabolic rate of species *i* relative to the time scale of the system and the non-dimensional constant *y*_*ij*_ is the ingestion rate of resource *i* by consumer *j* relative to the metabolic rate of *i*. The function *G*_*i*_(*B*) is the normalized growth rate of producer *i*, which follows logistic growth, 1 – *B*_*i*_/*K*_*i*_. The parameter *B*_*0*_ is the half saturation density. The efficiency *e*_*ji*_ is the fraction of the biomass of resource *i* consumed by consumer *j*, that is assimilated. We assumed efficiency *e*_*ji*_ = 0.45 for consumption of producers and *e*_*ji*_ = 0.85 for consumption of non-producers (Yodzis and Innes, 1992). We report results for systems in which *B*_*0*_ = 0.25 and the system-wide carrying capacity *K* = 5. The initial biomass of each species was uniformly drawn from [0.01, 0.1] for all simulations.

Simulations were run for 5000 model time steps at which point a top generalist predator invaded the food web. We assumed that the predator was an efficient generalist, with a fixed body mass consistent with a large secondary consumer (*M*_*predato*__r_ = Z^2.5^) and a scaled attack rate twice that of other species in the system. We note that the augmented predator attack rate is still within the range of empirically observed values (Peters, 1983). We used a simple rule for establishing the invading predator's feeding links upon invasion, where for each simulation the predator had a probability of 0.5 of establishing a feeding link with any consumer already present in the web. Consumers were explicitly defined as species whose shortest path along the network to any producer *P* = 1; the invader can consume herbivores or omnivores that are already present in the web, but not producers or other top predators. Following the invasion, each system was run for a further 5000 time steps. Cascade strengths were measured as *log*_*10*_ response ratios *log*_*10*_*B*_*post*_/*B*_*pre*_, where *B*_*post*_ and *B*_*pre*_ are aggregate producer community biomasses summed over all *n*_*R*_ producers and averaged over the final 100 time steps after and before predator invasions respectively. Biomasses were averaged in order to measure cascades for systems with oscillatory behavior in the steady state, which occurred in some of our simulations. Likewise, consumer level effects were calculated as *log*_*10*_ response ratios of aggregate consumer biomass. To ensure predators were not entering webs in which many species had gone extinct prior to their arrival, we set a limit on the maximum allowable number of extinctions prior to invasions at two, using *B*_*i*_ < 1 × 10^−15^ as the extinction threshold. In the event of an extinction before predator arrival, we allowed the extinct taxa to reinvade the system at an initial biomass equal to the extinction threshold. Numerical integration of ordinary differential equations was accomplished using the *deSolve*package in R (R Core Team, 2015).

To study whether features of the initial network structure were related to the response of systems to invading predators, we calculated associations between the cascade strengths and a suite of common network properties (Vermaat et al, 2009) using *ANOVA*. The properties we considered were connectance, species richness, characteristic path length, the fraction of species that are basal, intermediate and omnivorous, clustering coefficient, mean maximum trophic similarity and Clauset-Newman-Moore modularity (Clauset et al, 2004). We note that the frequentist statistical tests employed here were not used to assess significance since p-values are determined by the number of simulations. Instead, we follow the suggestion of White et al (2014) and use *ANOVA* as a framework for partitioning effect sizes and variance in these simulations and comparing effect sizes among covariates. We refer to these effects below using the notation *β*_*variable*_ where for instance *β*_*C*_is the connectance effect, which reflects the per unit impact of scaled *C* on the strength of cascades. Covariates were rescaled according to Gelman (2008) prior to analyses, to facilitate comparisons of estimated effects between different predictors that are necessarily on different scales.

Finally, we sought to understand the mechanisms underlying weak trophic cascades, as these cascades would be least likely detected in empirical studies. We operationally defined *weak cascades* as a less than twofold change in aggregate producer biomass after predator invasions. Under this definition, the mean cascade strength observed in terrestrial systems reported by Shurin et al (2002) would be considered weak (mean non-significant change by a factor of 1.1) whereas the average cascade strength reported for aquatic systems would be considered strong (mean change by a factor of 4.6). One possibility is that weak cascades are caused by diffuse predator effects (*sensu* Yodzis, 2000), whereby predator consumption is spread over multiple resources leading to overall weak population responses. In this scenario, species in each lower trophic level change only slightly in the same direction, and large community level biomass responses fail to emerge. Alternatively, weak cascades could occur in the presence of major changes to population biomasses if changes in strongly depressed species are offset by compensatory changes in the opposite direction by others (i.e., biomass compensation; Gonzalez and Loreau, 2009) in the producer or consumer guilds. To quantitatively assess these possibilities, we present a measure *µ* that quantifies the degree of biomass compensation among populations *i* in a trophic guild as

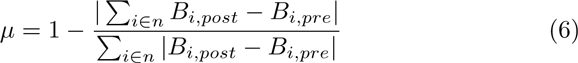

where the sum is over all *n* species in a trophic guild (e.g., all producers). This metric *µ* varies from 0 to 1, with 0 indicating that all species within a guild changed in the same direction (the biomass of all populations increased or decreased) and 1 indicating perfect compensation. If weak trophic cascades are typically accompanied by small *µ* values, then we conclude that weak cascades usually occur because top down effects are too diffuse to effect strong changes in individual producer populations and therefore aggregate producer biomass. Conversely, if weak cascades are typically accompanied by large *µ* then we conclude that compensatory changes by species in the same guild lead to a small net changes in aggregate biomass. Herein, we refer to compensation in producer and consumer guilds as *µ*_*R*_ and *µ*_*N*_.

## 3 Results

Predator invasions had moderate effects on aggregate producer biomass in most food webs ((Fig. 1). Producers changed by a factor of 1.7 on average across all simulations, and twofold changes in producer biomass occurred in only 31% of webs. Predator facilitation of producers was strongest in low richness and low connectance webs ((Fig. 2;*β*_*S*_ = –0.111, *β*_*C*_ = –0.012). Cascade strengths were also associated with other topological properties used to describe web structure (Vermaat et al, 2009). The strongest associations were observed between producer response ratios and species richness *S*, the fraction of basal species, the fraction of intermediate species and mean maximum trophic similarity (Table 1).

**Fig. 1.**
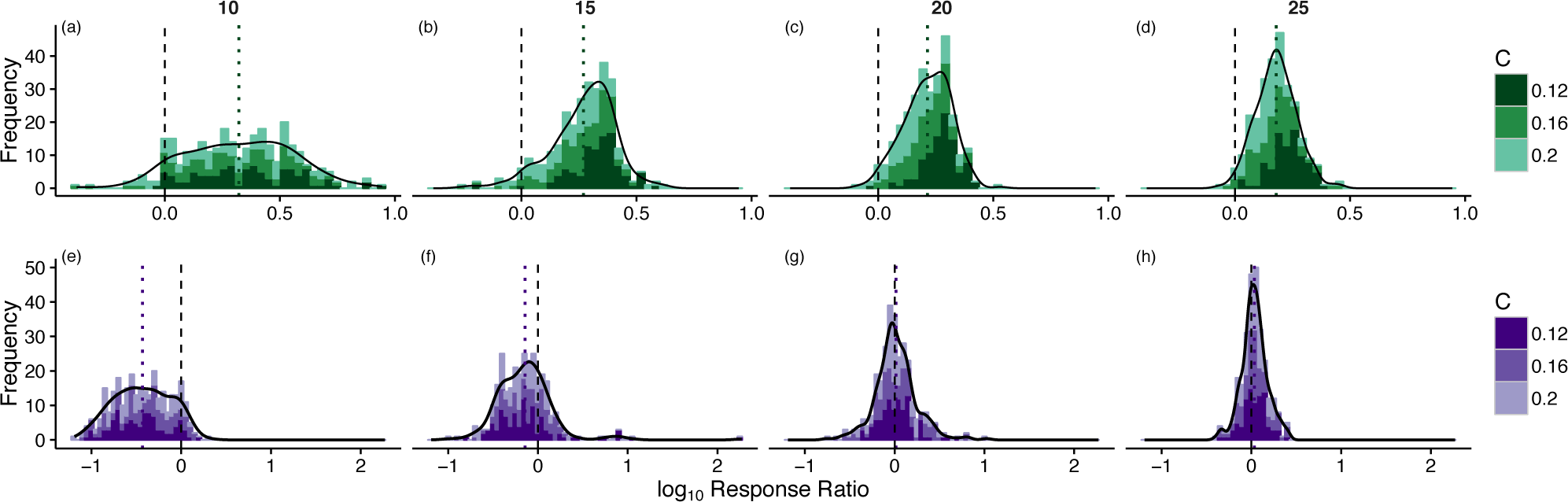
Stacked histograms of producer cascade strength frequency distributions for webs of different richness (panel columns) and connectance (green shading) values. The green dotted lines mark mean cascade strengths for reference. (e–h) Consumer cascade strength frequency distributions for webs of different richness (panel columns) and connectance (purple shading) values. The purple dotted lines mark mean consumer cascade strengths for reference. Density estimation was accomplished using a Gaussian kernel.

**Fig. 2.**
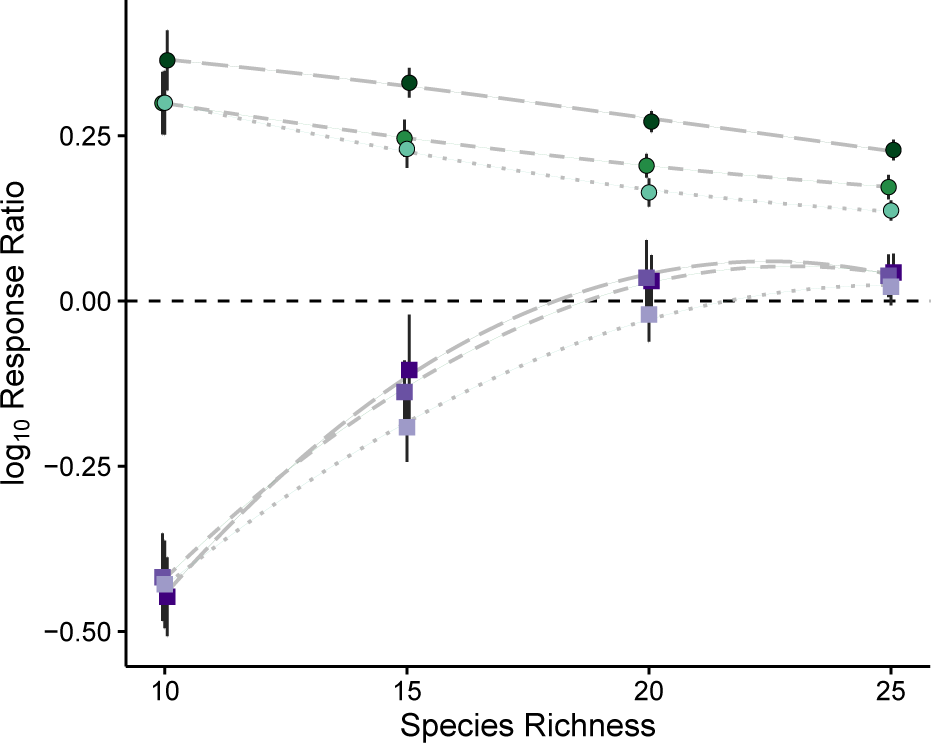
Relationships between species richness *S*, connectance *C* and cascade strengths in the producer (green circles) and consumer (purple squares) guilds. Points and error bars represent mean cascade strength ± 2 *SEM* and lines show results of *loess* regression to raw simulated data. Colors are the same as in Fig. 1. Dotted, short and long dashed lines correspond to webs with connectance values of 0.12, 0.16 and 0.2 respectively.

**Table 1.** Results of *ANOVA. β* indicates the estimated regression coefficient.

| Guild | Food Web Property | $\beta$ |
| --- | --- | --- |
| *Producers* |  |  |
|  | Species Richness | -0.111 |
|  | Connectance | -0.012 |
|  | Char. Path Len. | 0.08 |
|  | Frac. B | 0.21 |
|  | Frac. I | -0.116 |
|  | Frac. Om | 0.001 |
|  | Modularity | 0.014 |
|  | Clustering Coef. | 0.009 |
|  | Mean Max. Similarity | -0.095 |
| *Consumers* |  |  |
|  | Species Richness | 0.741 |
|  | Connectance | 0.156 |
|  | Char. Path Len. | 0.002 |
|  | Frac. B | 0.218 |
|  | Frac. I | -0.084 |
|  | Frac. Om | 0.014 |
|  | Modularity | 0.077 |
|  | Clustering Coef. | -0.084 |
|  | Mean Max. Similarity | 0.038 |

The magnitudes of consumer response ratios were more strongly correlated with most food web properties (Table 1), suggesting that the sensitivity of a guild’s log response ratio to initial network conditions may depend on trophic position; topology appears to exhibit relatively strong associations with changes in consumer level biomass following predator invasions compared to lower trophic levels. Depression of consumer biomass by the predator was strongest in low richness and weakly connected webs ((Fig. 2; *β*_*S*_ = –0.741, *β*_*C*_ = –0.156) with fewer basal species and less modular, more clustered network configurations (Table 1).

Producer compensation *µ*_*R*_ was negatively correlated with cascade strengths across all simulations ((Fig. 3a; Pearson’s, *r* = –0.34), suggesting that compensation among producers was in part responsible for masking cascades at the producer community scale (e.g., compare (Figs. 3b & 3c). This result is recapitulated by the high frequency of simulations characterized by stronger trophic cascades and almost no producer compensation ((Fig. 3a, dark shaded area). Indeed, of the webs that exhibited weak producer cascades (i.e., aggregate producer biomass increased by less than a factor of 2), 90% contained at least one producer population that more than doubled despite a weak community scale cascade. Taken together this suggests that weak cascades were in large part caused by producer compensation, leading to a small net changes in aggregate biomass. However, the magnitude of compensation was weakly correlated with other topological food web properties (*Supplementary Materials*), suggesting that predicting compensation at the scale of the trophic guild will require more detailed information than simple topological descriptors of ecological network structure.

**Fig. 3.**
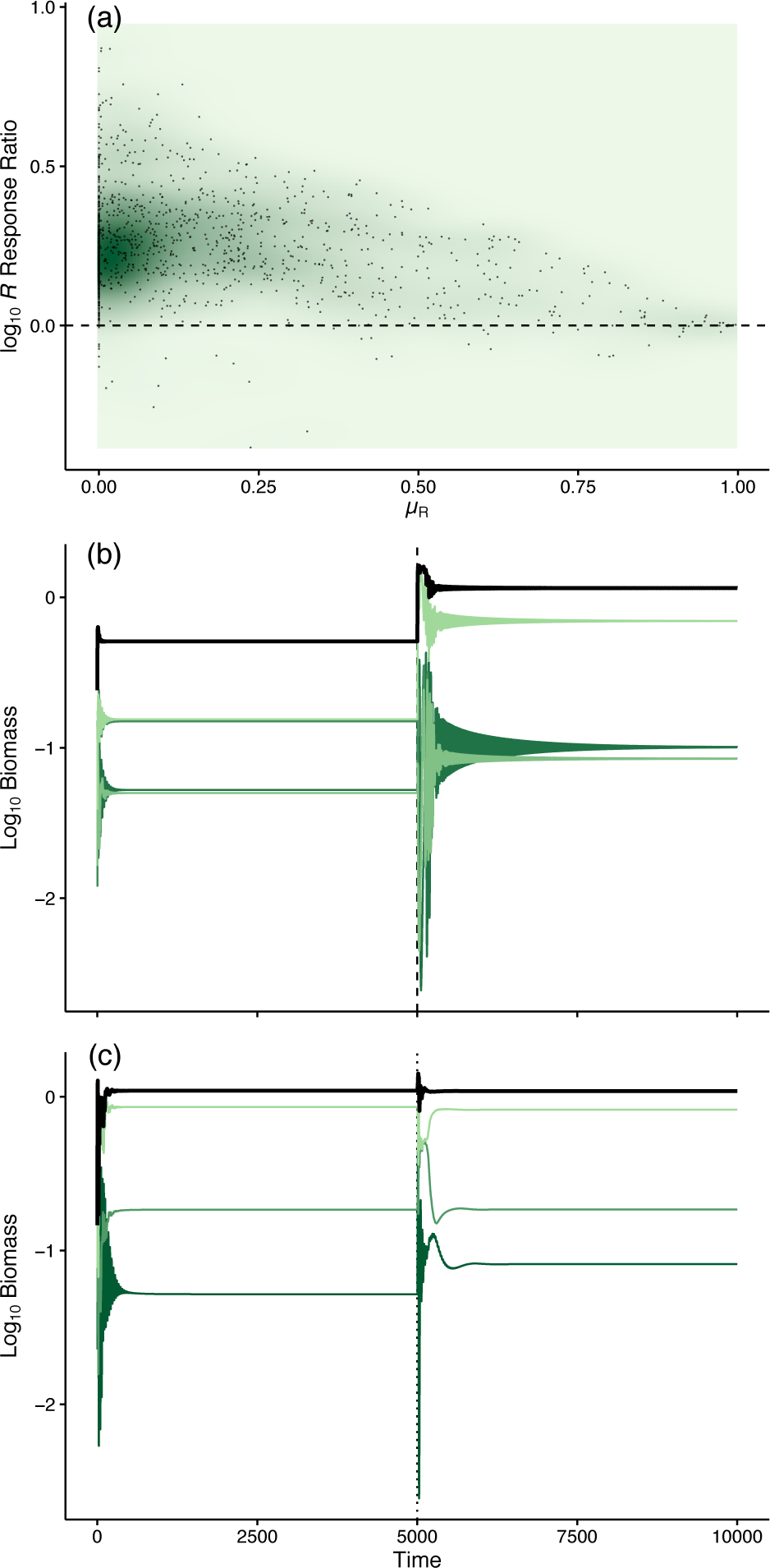
(a) Scatterplot showing the negative relationship between the producer cascade strengths and producer compensation, *µ*_*R*_. Points represent individual simulations. The background is shaded according to a 2D Gaussian kernel used for density estimation, where darker shades represent denser regions. A high density of stronger cascades with near-zero producer compensation is visible. (b) Example of a relatively strong cascade where compensation is weak. Colored green lines represent individual producer populations and the thick black line is aggregate producer biomass. A dashed line marks the predator invasion. (c) Example of a weak cascade arising from producer compensation.

Compensation in the consumer guild increased with species richness *S* and connectance *C* ((Fig. 4), explaining the shift in consumer effect size distributions toward zero visible in Figs. 1e-h. This suggests that two separate compensation mechanisms could explain weak cascades in webs. The first occurred more frequently in low richness webs, when strong depression of consumers cascaded to producer populations but failed to manifest at the guild scale because changes in some populations were offset by others in the opposite direction (i.e., *producer* compensation). The second occurred primarily in species rich webs ((Fig. 4), when top-down predator effects immediately diminished within the consumer guild due to *consumer*compensation. The strongest cascades occurred when both producer and consumer compensation was weak, which was most likely in low richness (lower *S*) and weakly connected (lower *C*) webs.

**Fig. 4.**
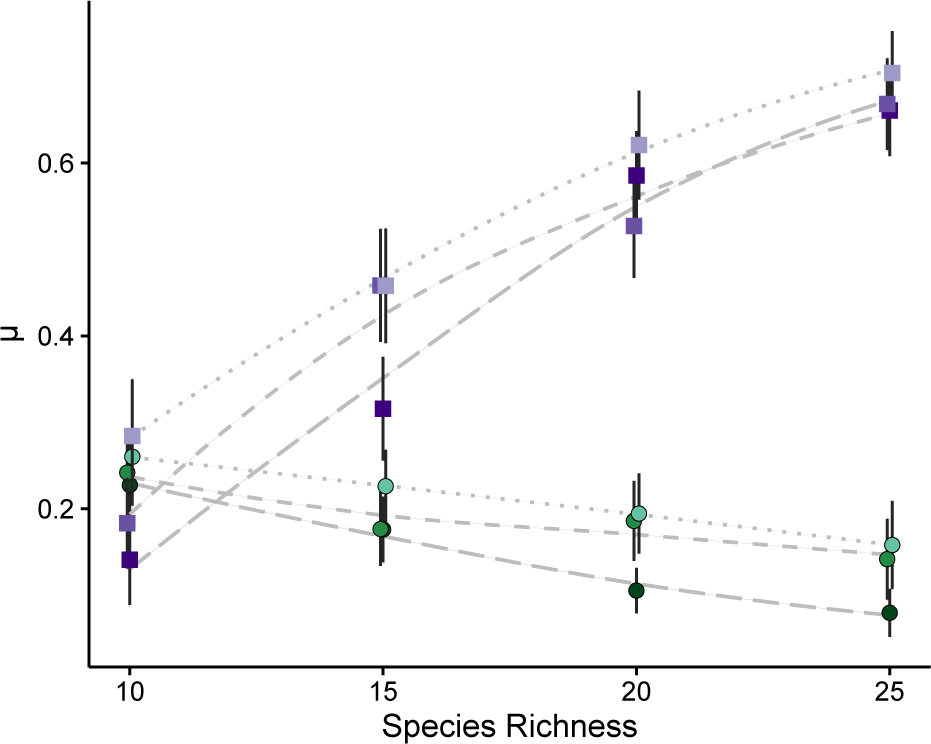
Relationships between species richness S, connectance *C* and compensation *n* in the producer (green circles) and consumer (purple squares) guilds. Points and error bars represent mean compensation ± 2 *SEM* and lines show results of *loess* regression to raw simulated data. Colors are the same as in (Fig. 1. Dotted, short and long dashed lines correspond to webs with connectance values of 0.12, 0.16 and 0.2 respectively.

## 4 Discussion

Our modeling study found that strong trophic cascades at the scale of the producer community are more likely to occur in weakly connected ecological communities with fewer species, a result that is in agreement with some previous interpretations of indirect effects and trophic cascades (MacArthur, 1955; Pace et al, 1999; Frank et al, 2006; Shurin et al, 2010). In most webs (90% of all simulations), at least one producer species doubled or more in biomass, yet strong guild scale cascades occurred in only thirty percent of simulations. Strong population level cascades were often offset by an opposite biomass change in other species so that the overall producer community biomass wasn’t strongly affected. Thus, restricting attention to trophic cascades as measured by changes in the overall biomass of a trophic guild makes it much less likely that the effects of an invading species will be detected. Strong top-down effects still occur in large and complex ecological networks, but observing them requires finer-grained observations than simply measuring total producer biomass (Polis et al, 2000). This is exemplified in high richness webs in particular ((Figs. 1d, 1h and (2), where changes in producer biomass occurred despite near-zero or slightly positive changes in aggregate consumer biomass on average. In almost all communities, the introduced top species had a strong effect on both the relative biomass of species and the dynamics of the community. Shifts in species composition due to compensation within a guild are more common than changes in overall community biomass, and may be a potentially potent indicator for species invasions (Schmitz, 2006).

Weaker cascades in large highly connected webs have been attributed to diffuse interactions among trophic levels in these systems (Leibold et al, 1997; Pace et al, 1999; Shurin et al, 2010). However, the observation that compensation frequently operated in multiple trophic guilds suggests a new hypothesis for the emergence of trophic cascades in complex food webs. Namely, changes at the top of webs have some chance of diminishing due to compensation within each trophic guild, as they cascade down to producers. If the network is structured in a way that precludes compensation from occurring in any of these guilds, then a strong cascade will emerge. Alternatively, top-down regulation has the capacity to diminish within a single trophic level if the propensity for compensation is high in that particular system, which can result from particular network architectures or exogenous abiotic forcing in real ecosystems (Gonzalez and Loreau, 2009). Notably, compensation was only weakly correlated with a suite of common topological food web descriptors, and thus additional research is needed to uncover the more nuanced features of food web architecture that drive compensatory responses at the scale of producer and consumer guilds. Experimental tests of the hypothesis discussed herein could be accomplished by adding conspecific generalist predators to replicate food webs with known topologies (e.g., experimentally assembled microcosms) and measuring them repeatedly through time. However, replicated food web experiments with repeated measures are scant and to our knowledge no such data exist to test the results presented here.

The present study looks at the role of increasing web richness and structural complexity on trophic cascades and the detection of the effects of species introductions. The model used, while more complex than those typically used in trophic cascade studies, is still highly idealized. The dynamics of real ecosystems often include many other non-trophic processes (Kéfi et al, 2015) which might dampen (or magnify) the cascading influence of top predators (Polis et al, 2000). One such example is that our study was restricted to models of closed systems. Evidence of cross-ecosystem cascades (Knight et al, 2005) and the effect of resource colonization rates on cascade strengths (Fahimipour and Anderson, 2015) suggest that extensions of our model to open systems will be a promising enterprise for further theoretical study. Future studies could build upon our model by exploring alternate assumptions and structures – for instance, other representations of primary production like fixed species-level *K* (e.g., Brose et al, 2006b), heterogeneity in resource productivity and edibility, different consumer functional responses, alternate assumptions about consumer metabolism and realistic ecosystem features such as detrital loops (e.g., Boit et al, 2012).

Identifying the abiotic and biotic features of ecosystems that regulate trophic cascades is a fundamental issue in ecology (Polis et al, 2000; Terborgh et al, 2010) and a practical problem for the management of invasive species, agricultural pests and zoonotic disease (Estes et al, 2011). While the present study identifies features of model food web architecture that influence cascades, the potential for compensation (Gonzalez and Loreau, 2009) which appears to be poorly predicted by ecological network structure, complex indirect interactions in real world ecosystems (Yodzis, 2000) together with insufficient data (Shurin et al, and issues of scale (Polis et al, 2000) combine to make the development of a predictive cascade theory of food webs a difficult problem.

## Acknowledgements

We thank Benjamin Baiser for useful discussions. This work resulted from the *Dynamics of and on Networks* workshop at the Santa Fe Institute in Santa Fe, NM. AKF was supported by the University of California Office of the President, the National Science Foundation under award no. 006741-002 and the Gordon & Betty Moore Foundation. KEA was supported by the University of California Academic Senate and the National Science Foundation under award no. DEB-1553718.

